# Alternative splicing redefines landscape of commonly mutated genes in acute myeloid leukemia

**DOI:** 10.1101/2020.05.21.107557

**Authors:** Osvaldo D. Rivera, Michael J. Mallory, Mathieu Quesnel-Vallières, David C. Schultz, Martin Carroll, Yoseph Barash, Sara Cherry, Kristen W. Lynch

## Abstract

Most genes associated with Acute Myeloid Leukemia (AML) are mutated in less than 10% of patients, suggesting alternative mechanisms for gene disruption contribute to this disease. Here we find a set of splicing events that disrupt the expression of a subset of AML-associated genes, including EZH2 and ZRSR2, independent of known somatic mutations. Most strikingly, in at least one cohort, aberrant splicing triples the number of patients with a reduction in functional EZH2 as compared to that predicted by somatic mutation of EZH2 alone. Together, these results demonstrate that classical mutation analysis underestimates the burden of functional gene disruption in AML and highlights the importance of assessing the contribution of alternative splicing to gene dysregulation in human disease.

---

Acute Myeloid Leukemia (AML) is an aggressive hematologic cancer in which malignant myeloid precursor cells impair hematopoiesis and induce bone marrow failure. AML is the second most common type of leukemia diagnosed in adults and, in spite of clinical efforts, the AML overall 5-year survival rate stands at roughly 25%^1,2^. One complicating factor in the prognosis and treatment of AML is that genomic lesions show complex patterns of cooperation and mutual exclusivity. Recurrent somatic mutations, truncations and small translocations occur in approximately 70 different genes that encode a variety of cellular functions^3,4^. Patient tumors typically contain mutations in several of these AML-associated genes; however, no single gene is mutated in more than ∼35% of all patients with most mutations occurring in less than 10% of patients^3,4^. Therefore, somatic mutations may underestimate the prevalence of disease associated with these genes.

Alternative pre-mRNA splicing is a pervasive cellular process that contributes to proteome complexity among higher eukaryotes and modulates patterns of gene expression that govern many cell fate decisions^5^. Importantly, changes in splicing patterns, such as differential inclusion or exclusion of an exon, often results in loss-of-function, dominant negative or altered protein activity^6^. Indeed, many cancers including AML can harbor mutations in splicing genes, suggesting that changes in splicing can drive cancer. Despite the demonstrated impact of alternative splicing and its potential association in AML, few studies of AML have explored the role of alternative splicing as an additional mechanism to somatic mutations by which gene function maybe altered in this disease.

As a first step towards addressing this gap, we investigated the splicing variability of genes that have been identified as commonly mutated, truncated or translocated in this disease, across two cohorts of AML patients. Strikingly we find that for several of these genes, including EZH2 and ZRSR2, alternative splicing markedly reduces protein expression in many AML patients independent of somatic mutations. Therefore, analysis of mutational status and gene expression level alone underestimates the breadth to which dysregulation of a particular gene is present in AML. In particular, accounting for alternative splicing more than triples the number of patients with reduced function of the epigenetic factor EZ2H compared to that estimated from analysis of somatic mutations in this gene. Together our analysis demonstrates the contribution of alternative splicing to gene dysregulation in AML and highlights the importance of quantitatively assessing this aspect of gene expression in determining the prevalence of gene disruption in disease.

## Results

To ask how splicing variation might contribute to gene dysregulation we first focused on 70 genes that have been identified as commonly mutated, truncated or translocated in greater than 1.5% of AML patients^3,4^, hereafter referred to as AML-associated genes. In our initial analysis we used polyA-selected RNA-Seq data from 29 in-house AML patient samples obtained from the Penn Stem Cell and Xenograft Core (aka PENN cohort, **Fig. 1, Table S1**). Samples in the PENN cohort were isolated by leukapheresis or from PBMCs (**Table S1**), and RNA-Seq data was generated from samples with an AML blast purity of greater than 90%. Thus, samples within the PENN datasets can be quantitatively compared without concern of confounding factors tied to sample purity. We obtained greater than 50 million paired-end reads for each of these 29 samples and quantified splicing patterns using the MAJIQ algorithm, which is optimized to detect and quantify complex and novel splicing events and thus is not limited to previously observed or expected splicing patterns^7^.

**Fig. 1.**
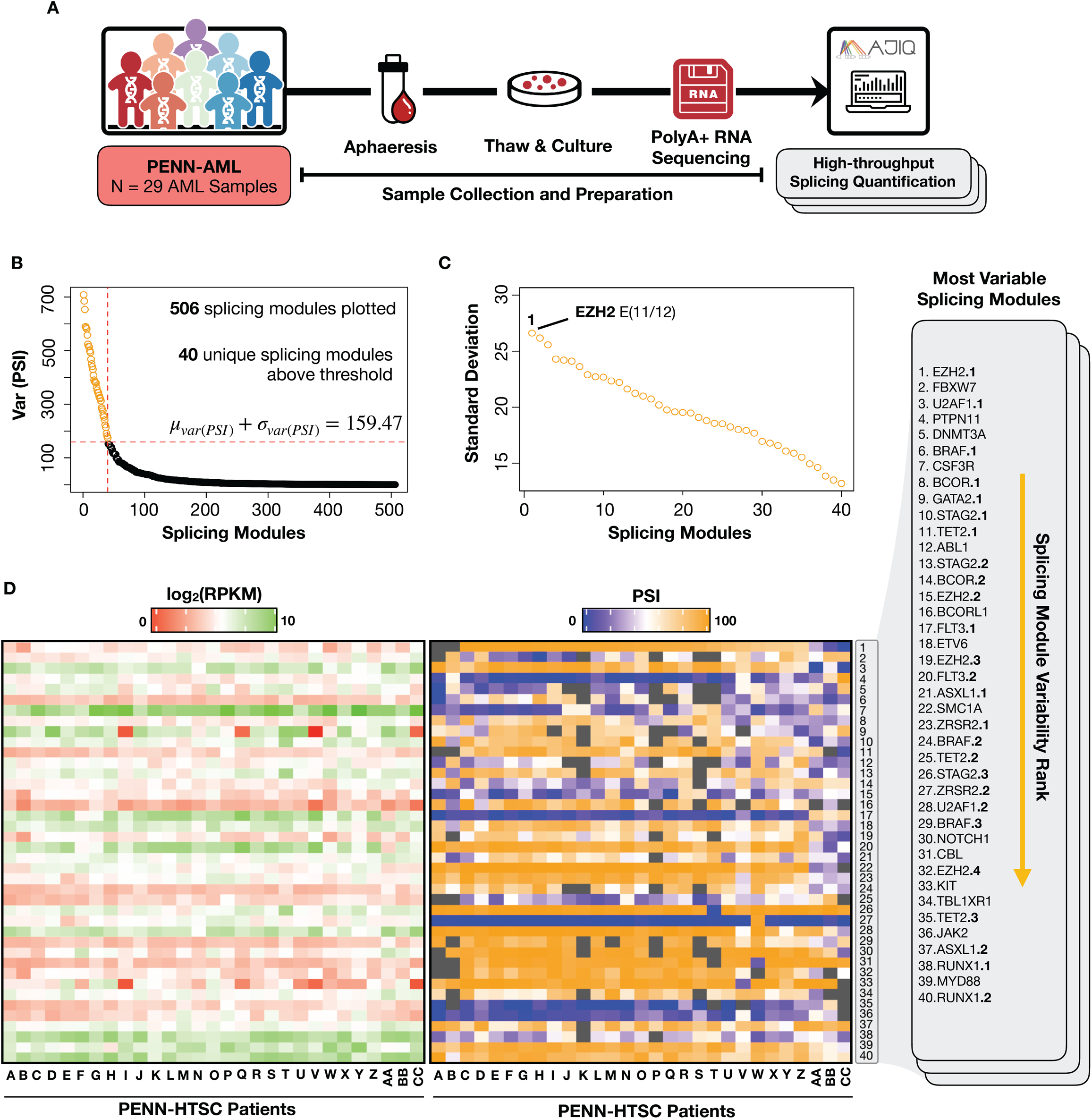
Splicing variability of AML-associated genes across the AML patient cohort. (**A**) Workflow of transcriptomic analysis from 29 primary AML samples obtained from the Penn Tumor Bank (**B**) Distribution plots of 506 splicing modules (x-axis) that were quantified in at least 80% of patients, sorted by the most variant PSI for each module (y-axis). The red cutoff line identifies modules with a variance one std dev (105.21) above the mean (54.26). (**C**) The 40 splicing modules above the threshold in panel B replotted according to standard deviation of PSI. The most variable module is indicated in the plot. See Figure 2 for details. (**D**) Heatmap showing transcript abundance values (left) and PSI (right) for each of the 40 highly variable modules. Rows are sorted by module # from panels B and C, with a list of these 40 modules on the far right. Where more than one variable module is observed within a gene, these are differentiated by a number after gene name. Columns in both heatmaps are identically ordered and sorted by based on PSI of the most variable module (EZH2 exon 11/12). Patients were then assigned letters consistent throughout this study. Gray indicates an unquantifiable module in a given sample.

Within the 70 AML-associated genes, we identified ∼500 splicing modules that are quantifiable in at least 80% of the PENN cohort (at least 24/29 patients, **Extended Dataset S1**). For this study, we define a splicing module as a gene interval (exons and introns) that can be spliced together in more than one pattern (e.g. see **Fig. 2A**). We are particularly interested in splicing variation that underlies patient heterogeneity and may be useful in stratifying patients. Therefore, we sorted these ∼500 modules by variance across patients (**Fig. 1B**). For each module we identified the most variable junction within the module and then calculated how much the percent usage of this junction (defined as Percent Spliced Isoform or PSI) deviates across the samples. Notably, this sorting identified 40 modules that exhibited a variance greater than one standard deviation above the mean variance for all modules (**Fig. 1B**). We also present these 40 “highly variable” modules based on their standard deviation of PSI (**Fig. 1C**) as well as the actual PSI for each patient (**Fig. 1D**, right heatmap). These 40 highly variable modules are within 25 of the 70AML-associated genes, with several genes harboring multiple highly variable modules (**Fig. 1D**). Thus, splicing variation in AML cells is common in transcripts of known leukemic oncogenes.

**Fig. 2.**
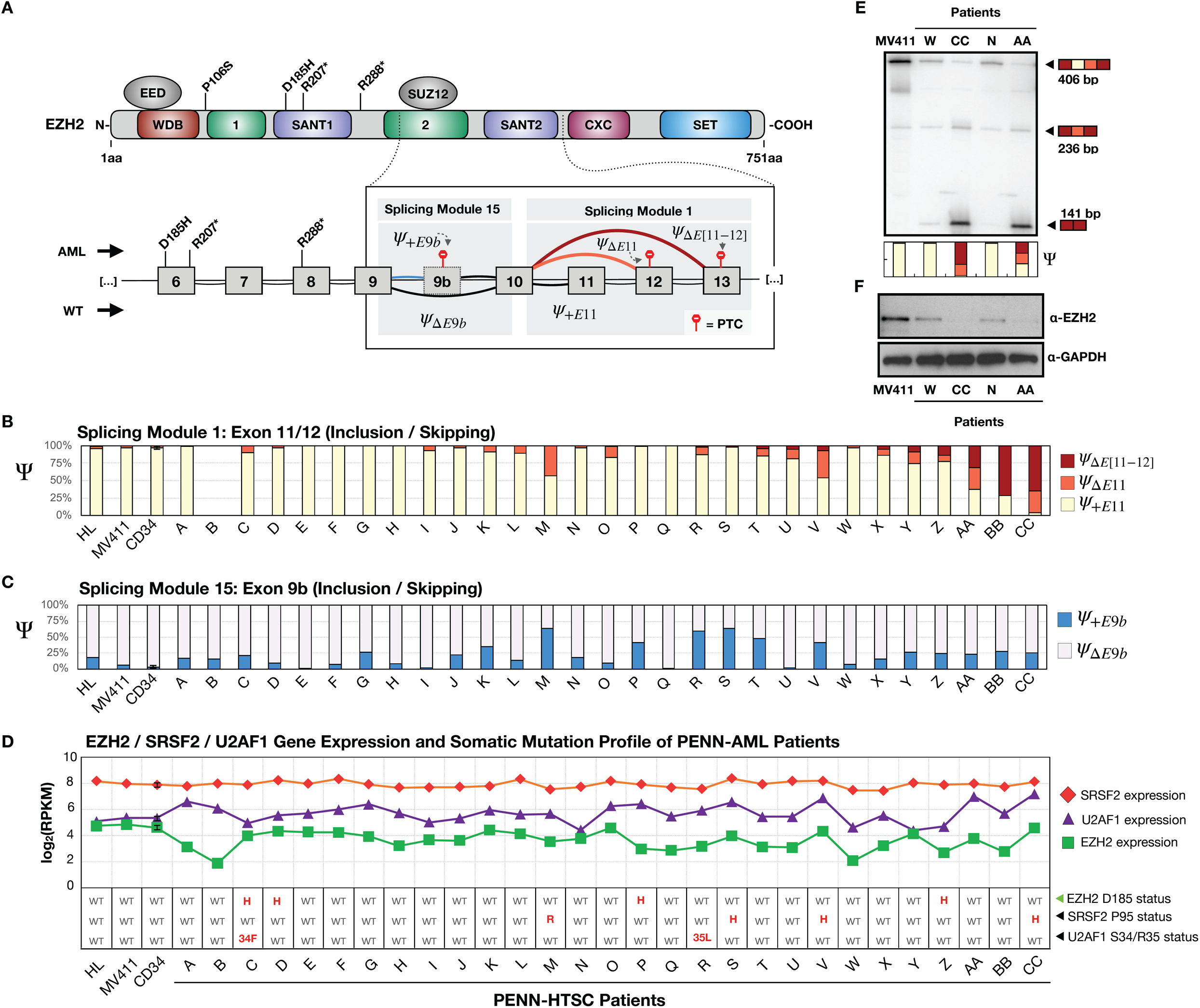
Splicing, expression and mutational analysis of EZH2 across the AML patient cohort. (**A**) Schematic of EZH2 protein domain structure and corresponding exon connectivity in EZH2 represented by the two most variable splicing modules in EZH2 from Figure 1. Also indicated are known AML related cis-mutations in EZH2 found in the PENN cohort (asterisk indicates nonsense mutations) and induced premature stop codons (PTC, red). (**B**) PSI values for exon 11 and 12 skipping in patients, cell lines (HL-60, MV411) and CD34+ normal cells (**C**) PSI values for exon 9b inclusion as in panel B. (**D**) gene expression (top) and mutational status (bottom) of EZH2 and two known regulatory proteins SRSF2 and U2AF1 in the samples corresponding to panels B and C above. (**E**) RT-PCR validation of splicing patterns of exons 11 and 12 from selected patients. (**F**) Western blot analysis of EZH2 expression from selected patients.

Gene expression for these same 70 genes showed overall less variability than splicing, and none of the genes harboring highly variable splicing modules were amongst those with highly variable expression (**Fig S1**). Moreover, for all of the 40 highly variable modules, splicing variability is independent of differences in transcript abundance, with the possible exception of modest correlation in the case of DNMT3A, ABL1 and STAG2.3 (**Fig. 1D, Table S2, Fig S2A**). We also determined whether there were correlations between PSI values and common, known pathogenic mutations within the same gene, and only observe significant correlation (□>|0.5|, p<0.01) for four of the highly variable modules (STAG2.3, TET2.1, SCM1A and KIT; **Table S3, Fig. S2B**). In these four cases, however, the correlation is driven by one or two patients. Therefore, none of the data strongly suggest that cis-mutations or expression drive splicing differences. Rather, we conclude that splicing variability represents differences in the transcriptome that are not captured by standard genetic and gene expression analysis in AML patients.

To understand how splicing variability may contribute to protein expression and function in patients, we first focused on EZH2 because it contains the most highly variable splicing module (**Fig. 1C, Fig. 2**). The EZH2 gene encodes the catalytic component of polycomb-repressive-complex-2 (PRC2) that confers di- and trimethylation on lysine 27 of histone H3 (H3K27me2/3)^8^. The most highly variable splicing event corresponds to skipping of exon 11 and/or 12 in the EZH2 gene (**Fig. 2A**, splicing module 1), which results in either case in a premature termination codon (PTC) prior to the catalytic SET domain. Skipping of these exons is not observed in two leukemia-related cell lines, MV411 and HL-60, and only minimally in normal CD34+ cells, but varies from 99% (e.g. patient CC) to 0% (patient A) in the PENN cohort (**Fig. 1D, 2B**). This complex exon skipping event has not previously been described, despite its prevalence in our and other AML data (see below), presumably because unlike the analysis done here, most splicing quantification methods focus on binary splicing choices and exclude events with more than two outcomes. Interestingly, variable splicing module 15 also corresponds to a PTC-inducing splicing event in EZH2, in this case through inclusion of variable exon 9b (**Fig. 1D, 2A, S3**). Inclusion of EZH2 exon 9b has previously been observed in some AML and MDS patients, and has been shown to induce nonsense-mediated decay (NMD) leading to loss of protein expression and a corresponding reduction in H3K27me3 levels^9^. Consistently, we observe over 40% inclusion in exon 9b in six of the PENN patients (**Fig. 2C**, patients M, P, R-T, V). However, there is no correlation between inclusion of exon 9b and splicing of exons 11/12 in EZH2 (**Fig. 2B,C**).

Importantly, we have validated skipping of EZH2 exons 11/12 in several of our AML patient cells (**Fig. 2E**) and observe the predicted decrease in EZH2 protein in these same samples (**Fig. 2F**). Heterozygous loss-of-function (LOF) somatic mutations in EZH2 have been described in AML but are only observed in a small percentage of patients ^3^. Consistently, we find that 4 of our 29 patients (14%) harbor a somatic mutation in EZH2 (**Fig. 2D, Table S3**). By contrast, we find 8 additional patients (28%) for which greater than 50% of their EZH2 transcripts fail to encode full-length protein due to skipping of exon 11/12 and/or inclusion of exon 9b (patients M, R-T, V, AA-CC). Therefore, taking splicing patterns into account in the transcriptomic analysis more than triples the percentage of recognized EZH2-deficient patients (14% to 42%), consistent with the notion that splicing variation is an underappreciated contributor to protein dysregulation in AML.

Previous studies have shown that one driver of EZH2 exon 9b inclusion is mutation of the splicing regulatory protein SRSF2. Specifically, mutation of SRSF2 proline 95 (P95) to histidine, lysine or arginine alters the RNA binding specificity of this splicing factor, sresulting in its binding to sequences within EZH2 exon 9b to promote exon 9b inclusion^9^. Using the RNA-Seq data we determined that, of the 6 patients in the PENN cohort that exhibit highest inclusion of EZH2 exon 9b, 2 carry the P95H mutation, while a third has the P95R mutation (**Fig. 2D**). A second mutation that has been shown to correlate with high inclusion of EZH2 exon 9b in MDS is a S34F mutation in the core splicing factor U2AF1^10^. We detect this mutation in patient C, which has low inclusion of exon 9b (**Fig. 2D**), however mutation in the neighboring residue in U2AF1(R35L) does co-exist with high exon 9b inclusion in patient R (**Fig. 2D**). U2AF1 R35L has been detected in AML^4^, but has not previously been shown to drive inclusion of EZH2 exon 9b. Notably no mutations in SRSF2, U2AF1 or EZH2 correlate with skipping of EZH2 exons 11/12, nor does this splicing event correlate with the gene expression level of EZH2, SRSF2 or U2AF1 (**Fig. 2D**). Therefore, none of the genomic features previously described as correlated with functional EZH2 expression in AML would predict the loss of EZH2 protein observed upon skipping of EZH2 exons 11 and/or 12. Moreover, at least one instance of high inclusion of EZH2 exon 9b lacks known mutations in SRSF2 or U2AF1 (patient T), while another patient (CC) has the SRSF2 P95H mutation but low inclusion of exon 9b, suggesting other yet-unidentified drivers of this splicing event.

Having identified unanticipated splicing variations in the PENN cohort that are informative regarding the expression and function of AML-associated genes, we next determined whether these variations are observed in larger cohorts of AML patients. We re-analyzed data from ∼440 AML patient samples published as part of the BEAT-AML project^11^. Overall, we find that splicing is generally more variable in both the PENN and BEAT-AML cohorts than is observed in normal CD34+ cells (**Fig. 3A**). Moreover, at least 30 of the 40 modules that exhibited a variance of greater than 160 in the PENN cohort (**Fig. 1B**), also had a variance above this threshold in the BEAT-AML samples (**Fig. 3B, S4**). By contrast, analysis of the same splicing modules in CD34+ cells from 17 healthy donors revealed much less variation (**Fig. 3B**). Therefore, the majority of the highly variable splicing events we detect in our in-house population is reflective of AML patients more generally. Even more remarkably, 23 of these splicing events (including EZH2.1) show a high degree of correlation in both the PENN and BEAT-AML cohorts, suggesting that these events are co-regulated with one another (**Fig. 3C**). This cluster of co-regulation between splicing modules is not observed in the normal CD34+ cells (**Fig. 3C**).

**Fig. 3.**
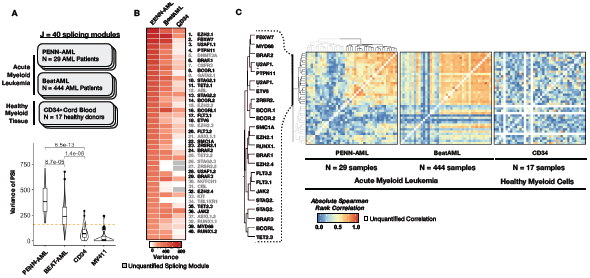
Splicing variations in the PENN cohort are also observed in additional AML cohorts. (**A**) Variance of PSI for the 40 highly variable splicing modules quantified in the PENN cohort from Figure 1, plotted also for that observed in 444 samples from the BEAT-AML study or 17 healthy CD34+ cell samples from Leucegene publicly available data. Orange dashed line indicates the threshold variability from Figure 1B. (**B**) Heat map of the variance in PSI observed in the BEAT-AML cohort and normal CD34+ control cells of the 40 most variable modules from the PENN cohort. Modules are named as in Figure 1D. Colors of modules are based on presence in black or grey clusters in panel C in this figure. (**C**) Spearman correlation of each of the variable modules with all others shown as a heatmap. Modules were clustered based on correlation values, highlighting a cluster of 23 highly correlated splicing modules (black bar, names listed on left). Clustering was done based on PENN data (left panel). The same ordering reveals strong clustering and correlation also in the BEAT-AML data (middle panel) but not in the CD34+ cells (right panel).

**Fig. 4.**
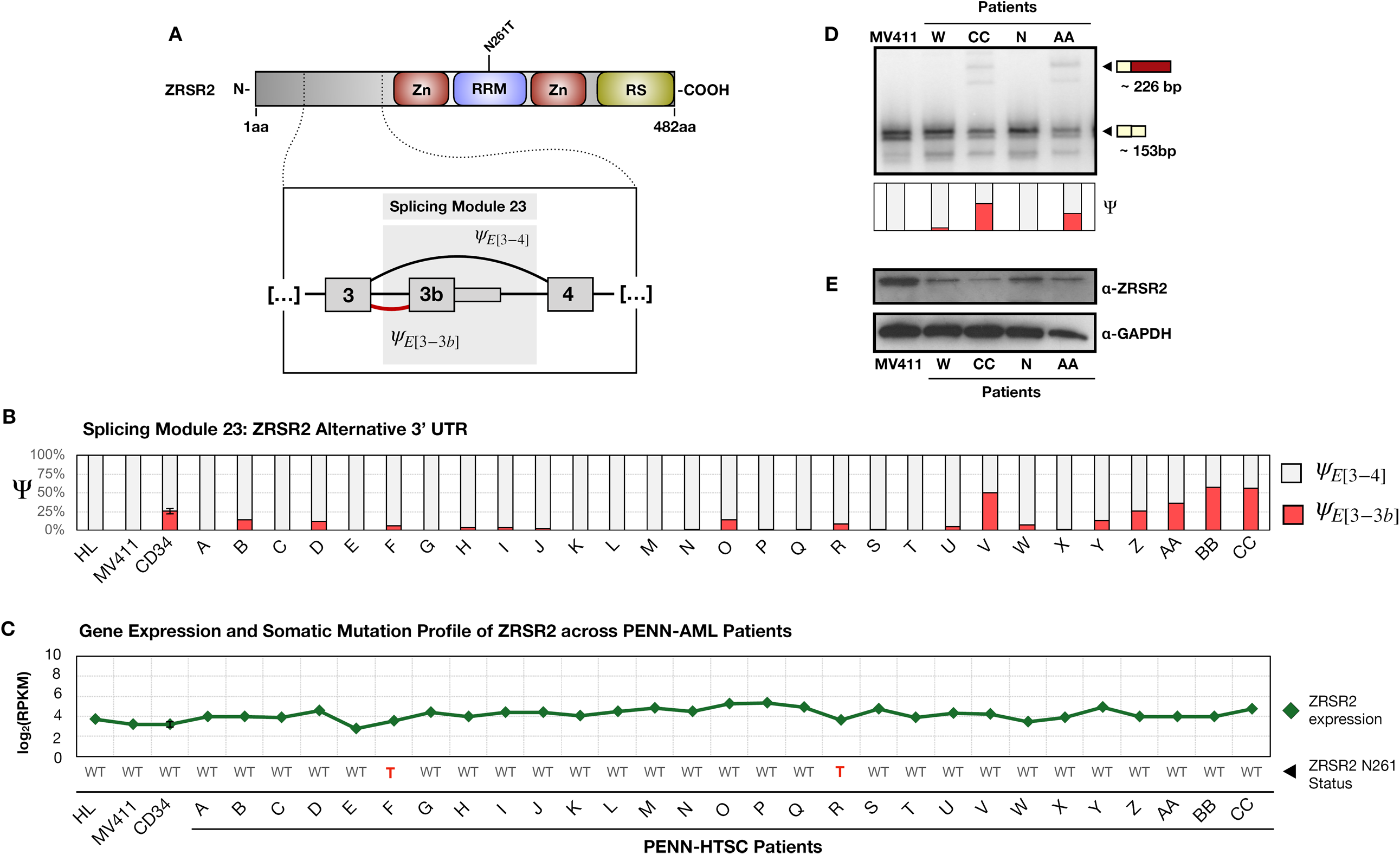
Splicing, expression and mutational analysis of ZRSR2 across the PENN AML patient cohort. (**A**) Schematic of ZRSR2 protein domain structure, and exon connectivity in ZRSR2.1, module 23 from Figure 1 and 3. (**B**) PSI values for exon 3 splicing to alternate exons 3b or 4 in PENN cohort patients, cell lines (HL-60, MV411) and CD34+ normal cells (**C**) Gene expression (top) and mutational status (bottom) of ZRSR2 in the samples corresponding to panel B. (**D**) RT-PCR validation of use of exons 3b and 4 from selected patients. (**E**) Western blot analysis of ZRSR2 expression from the same patients as panel D.

We note that among the genes that contain the co-regulated splicing events are U2AF1 and ZRSR2, both splicing factors themselves that participate in the recognition and use of 3’ splice sites^5,12,13^. Notably, AML-associated mutations have been shown to impact the specificity of RNA binding of U2AF1 or result in loss of function of ZRSR2^14,15^. Given that changes in the expression or function of either of these proteins are predicted to impact the splicing of other genes in AML we were particularly interested in whether the splicing variation we observe in these genes impact the encoded protein. The most highly variable splicing module in U2AF1 involves the inclusion of both exon 3 and 3’, two tandemly duplicated exons (**Fig. S5A,B**, splicing module 3). Inclusion of either exon results in functionally active protein; however, if both exons are included, the reading frame is altered in exon 3’ to generate a stop codon. While we can confirm the inclusion of both exon 3 and 3’ by RT-PCR (**Fig. S5C**), this event represents less than 25% of the transcripts and thus, not surprisingly, does not correlate with a detectable decrease in full length U2AF1 protein (**Fig. S5D**). We also note the presence of another splicing module (U2AF1.2, module 28) that involves skipping of exon 5, which is predicted to result in an in-frame deletion of a portion of the UHM domain that dimerizes with U2AF2 (**Fig. S5A,E**); however, as we don’t detect any smaller U2AF1 protein in our Western blot (**Fig. S5D**) and U2AF1 transcript expression is high in patients that skip exon 5 (**Fig. S5F**), we cannot make any predictive statements about this isoform.

In contrast to the lack of detectable impact of U2AF1 variable splicing on protein expression, we do observe a clear reduction of full-length ZRSR2 protein that correlates with variation of splicing of module ZRSR2.1 (module 23, **Fig. 5**). In particular, 4 patients (V, AA, BB and CC) exhibit greater than 25% splicing from exon 3 to an alternative terminal exon 3b, rather than to exon 4 and beyond to encode the full-length protein (**Fig. 5A,B**). We also detect near-complete skipping of ZRSR2 exon 9 in one patient (module 27, patient W, **Fig. S6**), which is predicted to introduce a premature stop codon. Consistently, in the patients for which we have protein samples, we observe a marked decrease in full-length ZRSR2 in patients that contain aberrant levels of these splicing events relative to control patients or MV411 cells, despite the fact that the overall expression (RPKM) for ZRSR2 is essentially the same in all samples (**Fig. 5C-E**).

We next asked whether the high variable splicing modules shared any features suggestive of a common regulatory mechanism. Motif analysis for the highly variable modules revealed enrichment of a purine-rich element in the exons and a U-rich element in introns (T-rich in the genomic signature) as compared to non-variable modules (**Fig S7A**). ZRSR2 does not have a defined binding motif, but it is possible that the U-rich motif reflects sequences in a 3’ splice site that confer ZRSR2 regulation. We also determined if expression of known purine-rich or U-rich binding proteins exhibited any correlation with the splicing of the highly variable modules. Interestingly, we find a strong correlation (**Fig S7B,C**) between expression of SRSF1, and most of the 23 co-regulated modules from **Fig 3** (i.e. cluster 1). A similar but weaker trend was also observed for hnRNP F, but not for any of the other purine-rich (hnRNP H, hnRNP A1, TRA2A, SRSF7) or U-rich (TDP43, TIA1) factors, or other splicing regulatory factors that are mutated in some AML patients (U2AF1, SRSF2). While we are unable to modulate expression of ZRSR2, SRSF1 or hnRNP F in patient cells, and as predicted knock down of these factors in MV411 cells had no impact on splicing of these 23 modules (data not shown), our data does suggest potential regulatory factors that may drive at least some of the variability in splicing across AML patients.

## Discussion

In sum, our analysis here demonstrates splicing as a major contributor to gene dysregulation in AML. In particular, we find that alternative splicing of EZH2 and ZRSR2 lead to a higher penetrance of loss of function of these genes in AML than has been previously recognized. More broadly, our findings highlight the critical importance of assessing splicing patterns, in addition to genetic mutation and altered expression, in determining the contribution of individual genes to disease phenotype. While we demonstrate this to be true here for AML, we predict that assessment of splicing will be similarly relevant to the study of gene dysregulation in most other diseases. In addition, our data suggest the merit of larger studies with clinically annotated datasets to determine if splicing patterns correlate with response to chemotherapy and/or survival.

## Acknowledgments

The authors wish to acknowledge the Stem Cell and Xenograft Core of the Perelman School of Medicine and the Hematologic Malignancies Translational Center of Excellence of the Abramson Cancer Center for obtaining the AML patient samples and Roberto Bonasio for EZH2 antibodies. We also thank the High-Throughput Screening Core, and the Penn Center for Precision Medicine.

## Author contributions

O.D.R. analyzed data on the 70 AML-associated genes from the PENN and BEAT-AML datasets, M.J.M. carried out RT-PCR and western blot validations, M.Q.V. assembled the BEAT-AML data and carried out the initial MAJIQ analysis on these data, D.C.S. cultured patient cells and isolated RNA for sequencing, Y.B. directed the computational analysis, S.C. and K.W.L. conceived and directed the project and, with all authors, wrote the manuscript.

## Data and materials availability

newly generated RNA-Seq data from the PENN cohort, HL60 and MV411 cells is available under accession GSE142514.

## Supplemental Figure Legends

**Fig. S1.**
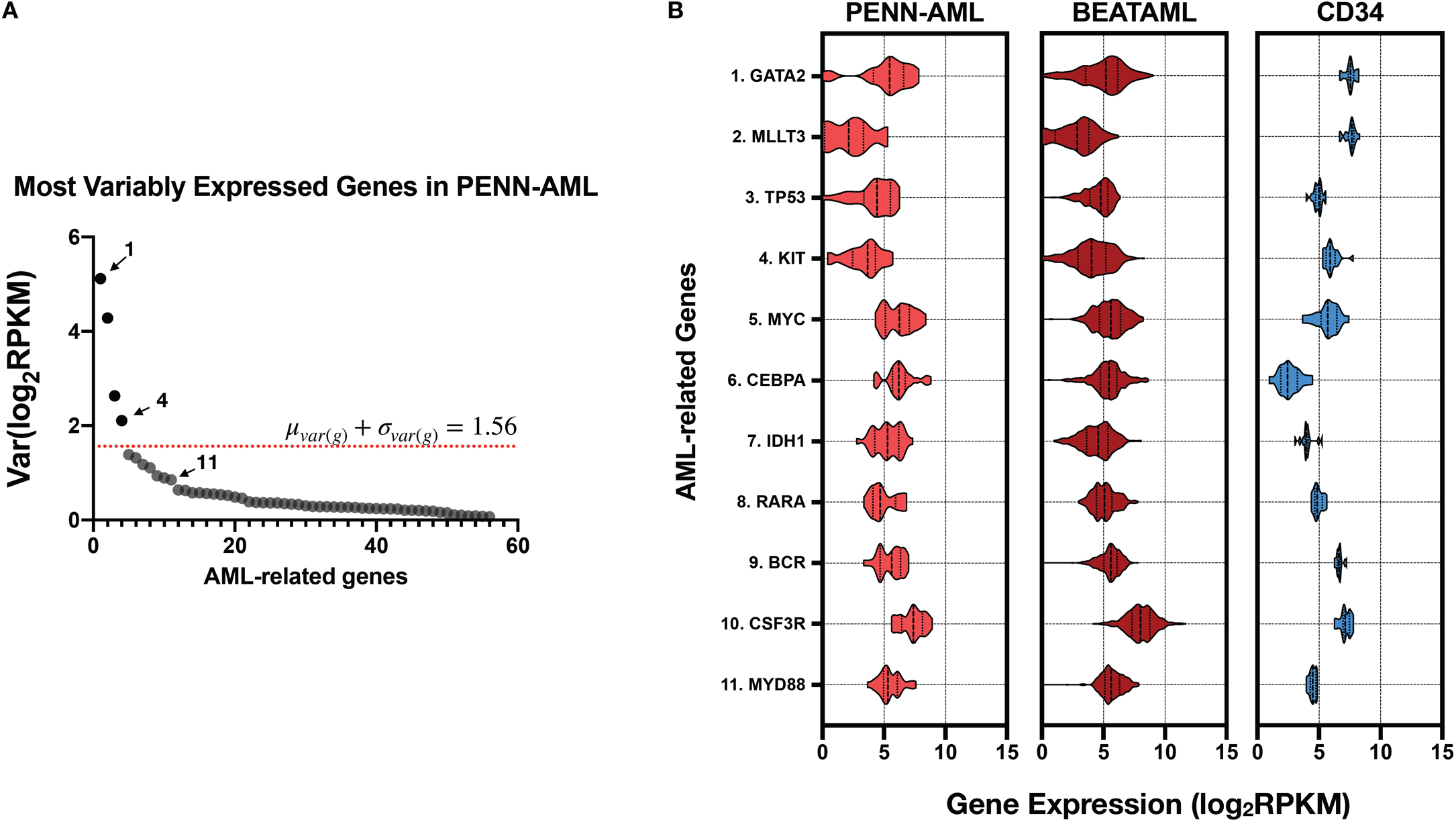
Genes with variable expression in AML are distinct from those exhibiting splicing variability **(A)** Distribution plot of variability in gene expression for the 70 AML-associated genes (x-axis) that were quantified in at least 80% of patients, sorted from most to least variable. The red cutoff line identifies modules with a variance one std dev (105) above the mean (54). (**B**) Distribution of expression values for the top 11 most variable genes in Penn and BEAT AML cohorts, plus 17 CD34+ cells from healthy donors.

**Fig. S2.**
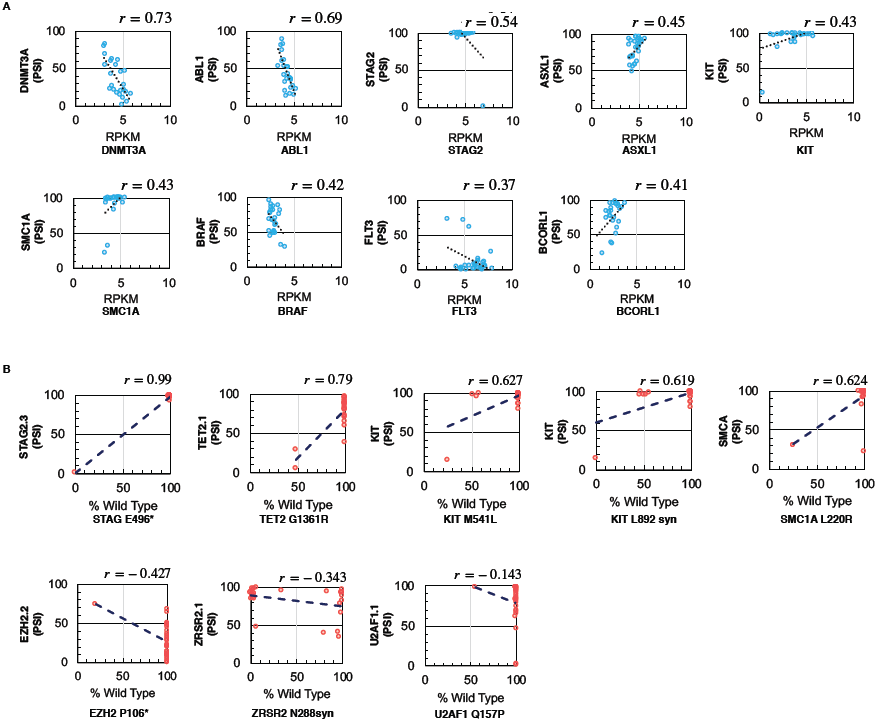
Splicing variability shows little correlation with cis-mutations **(A)** Correlation plots of RPKM vs. splicing variability in the same gene (PSI values) for splicing modules that exhibit some correlation with cis-mutations (top three) or splicing modules included in other figures in the study (bottom three). In all plots the Pearson r value is given. Further details are in Table S2. **(B)** Correlation plots of splicing variability (PSI values) vs. genetic mutations for splicing modules that exhibit some correlation with cis-mutations (top four) or splicing modules included in other figures in the study (bottom three). In all plots the Pearson r value is given. Further details are in Table S3.

**Fig. S3.**
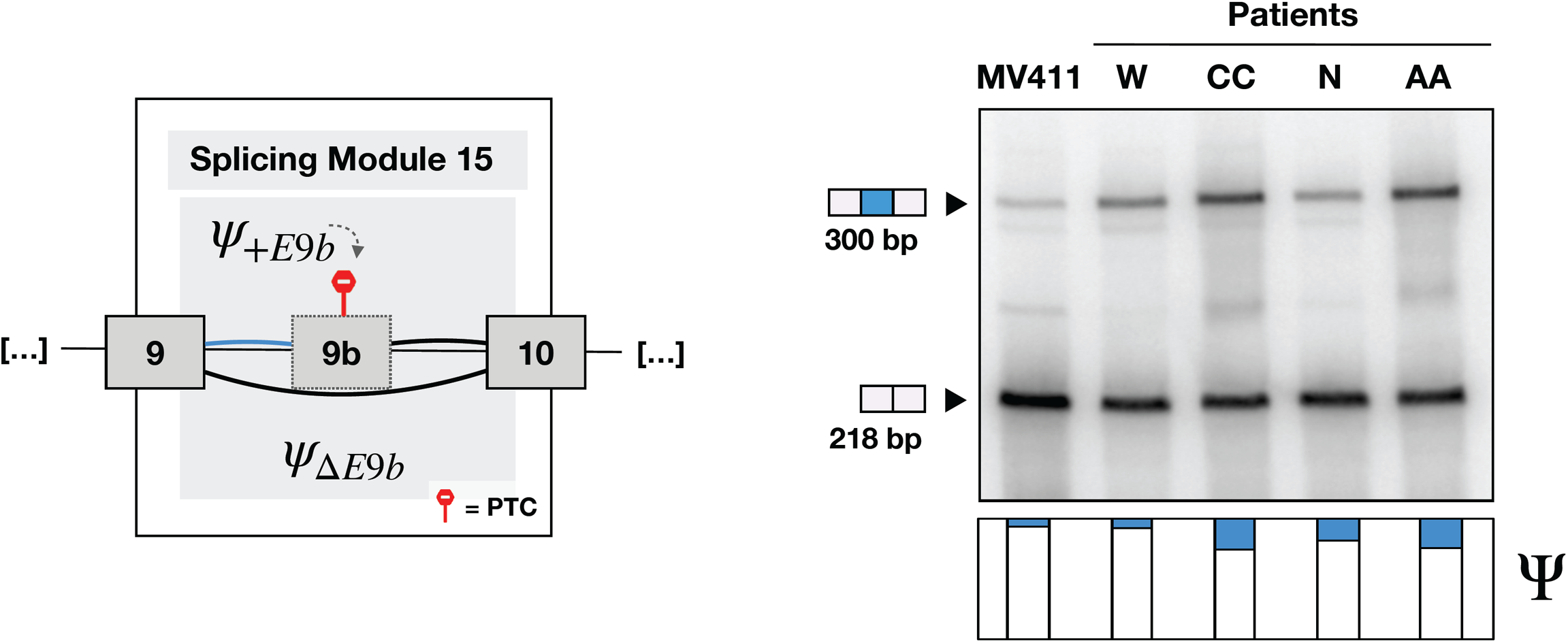
RT-PCR analysis of inclusion of EZH2 eon 9b (top) and the PSI values calculated from the RNA-Seq (bottom) from four representative patients. See also Figure 2.

**Fig. S4.**
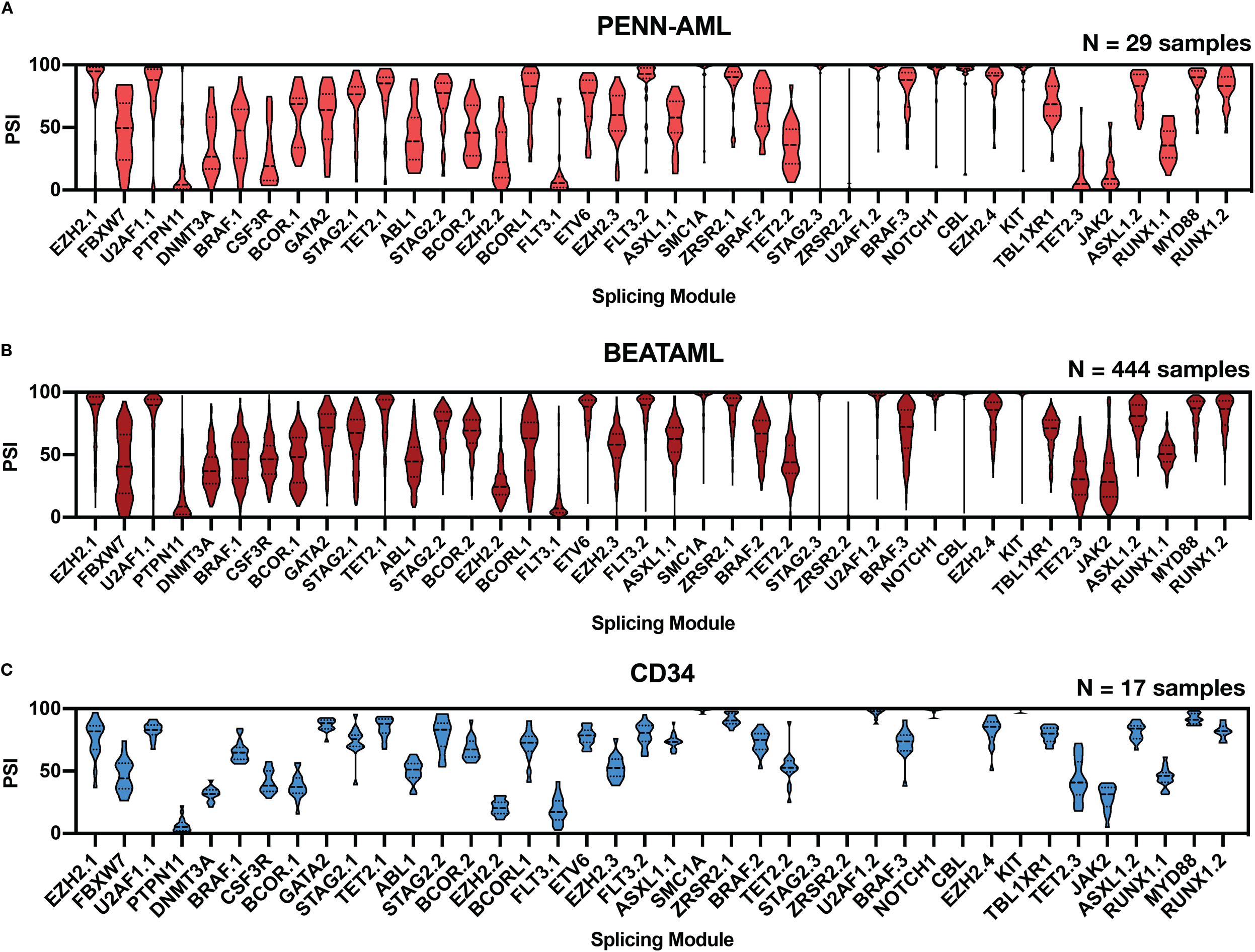
Distribution of PSI for the 40 modules from Figure 1 as observed in the BEAT-AML patient cohort and CD34+ cells from 17 healthy donors. See also Figure 3 and Extended Dataset S1.

**Fig. S5.**
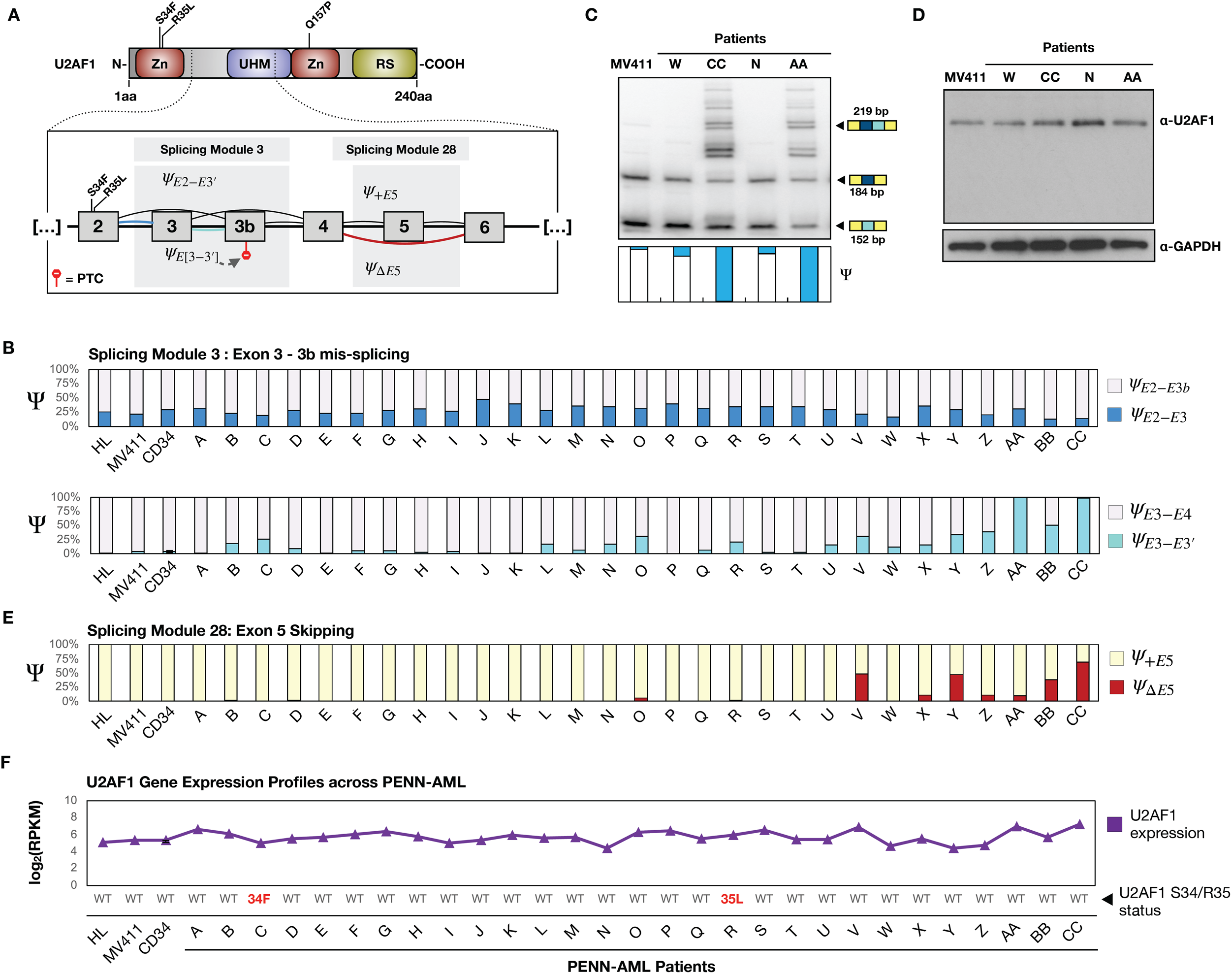
Splicing, expression and mutational analysis of U2AF1 across the PENN AML patient cohort. (**A**) Schematic of the U2AF1 protein domain structure, and exon connectivity in U2AF1, modules 3 and 28 from Figure 1 and 3. (**B**) PSI values for module 3; exon 2 splicing to alternate exons 3 or 3b or 4 (top) or 3 to 3b (bottom) in PENN cohort patients, cell lines (HL-60, MV411) and CD34+ normal cells. (**C**) RT-PCR analysis of module 3 splicing from selected patients. (**D**) Western blot analysis of U2AF1 expression from the same patients as panel C. (**E**) PSI values for exon 5 skipping (module 28) in PENN cohort patients, cell lines (HL-60, MV411) and CD34+ normal cells. (**F**) Gene expression (top) and mutational status (bottom) of U2AF1 in the samples corresponding to panel B and E.

**Fig. S6.**
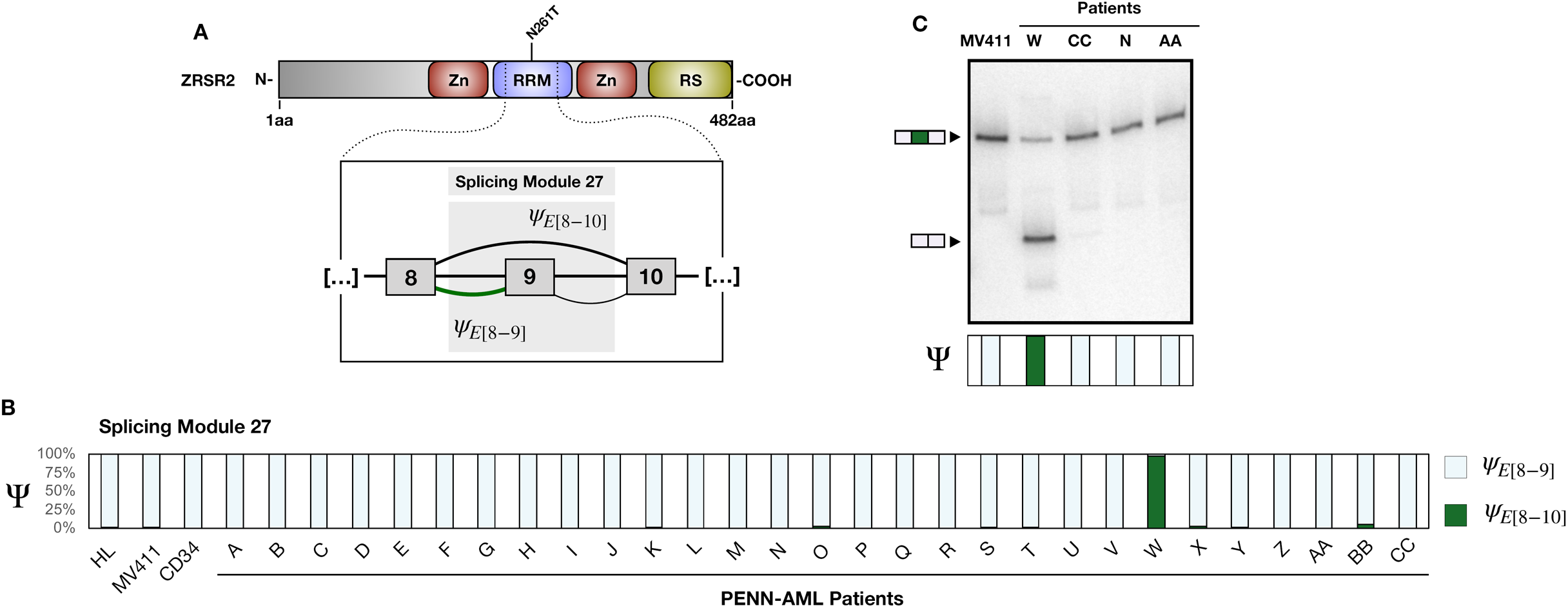
Splicing, expression and mutational analysis of ZRSR2 across the PENN AML patient cohort. (**A**) Schematic of ZRSR2 protein domain structure, and exon connectivity in ZRSR2.2 (module 27, exon 9 skipping). (**B**) PSI values for exon 9 skipping in PENN cohort patients, cell lines (HL-60, MV411) and CD34+ normal cells (**C**) RT-PCR analysis of exon 9 splicing from selected patients.

**Fig. S7.**
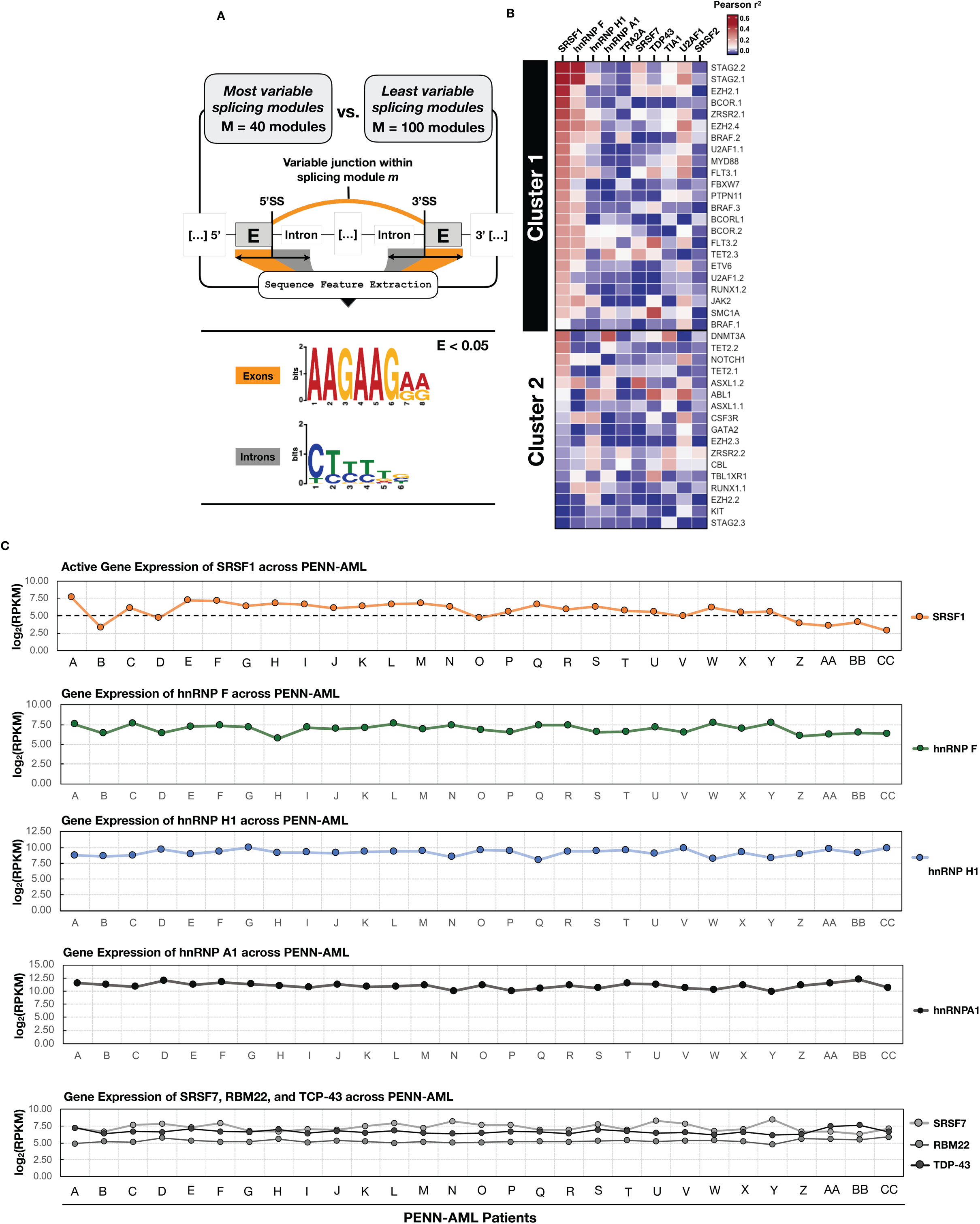
Motifs surrounding variable exons suggest mechanisms of regulation. (**A**) Motif enrichment analysis of exonic and intronic sequences encompassed in the high variable splicing modules. (**B**) Heatmap of correlation between expression of RBPs known to bind to GA- or U-rich sequences and PSI of highly variable splicing modules. (**C**) Detailed expression data for several RBPs from panel B.

## Methods

### AML Patient Samples

To construct the PENN dataset, AML blasts were first isolated from AML patients via size-exclusion centrifugation apheresis or from peripheral blood on ficoll gradiants and stored in the Penn Stem Cell and Xenograft Core at the University of Pennsylvania. To promote the molecular characterization of a homogenous cell population, AML patient samples were confirmed to be at least 90% blasts. CD34 cells from PENN were isolated from adult bone marrow. Total RNA was isolated from AML blasts by Trizol followed by DNAse treatment and reprecipitation. Poly(A) selection, library preparation, and paired-end sequencing was performed by GeneWiz.

### Poly-A mRNA Sequencing and Data Processing

High-throughput mRNA Sequencing files (.fastq) were pre-processed by Trim Galore (https://github.com/FelixKrueger/TrimGalore) to trim short-read adapter sequences from sample reads. Significantly larger datasets, such as the Beat-AML cohort were pre-processed with BBDuk (Decontamination Using Kmers, https://jgi.doe.gov/data-and-tools/bbtools/). Pre-processed reads were then aligned to the human genome (GRCh38 Genome Reference Consortium Human Reference 38, GCA_000001405.15) using STAR version 2.5.4b (https://github.com/alexdobin/STAR). The read-depth of RNA-Seq data from the Penn cohort ranged from 50 to 200 million reads per sample, with read lengths ranging from 75-150 nucleotides.

### mRNA Splicing Quantification and Alternative Splicing Detection

RNA-seq reads mapping to 70 AML-related genes were used for downstream splicing quantification analyses. The MAJIQ ^7^ splicing quantification tool (https://majiq.biociphers.org/) was used to identify all local splicing variations (LSVs) within the defined set of 70 AML-related genes across the PENN-HTSC cohort. An LSV is defined as a single exon alternatively joined to more than one RNA segment. Junctions involved in an LSV are quantified by Percent Splicing Index (PSI) values that represent the relative fraction of poly-A selected mRNA transcripts in which the aforementioned exon is joined to each of the alternative RNA segments. Variations in junction PSI represent the occurrence of a particular splicing pattern across RNA-seq data. To summarize our data, the quantified LSVs were used to construct *splicing modules* which model the splicing events that are occurring within AML-related gene transcripts. For every splicing module, the variance in PSI for junction j, or var(PSI_j_), across a particular cohort was calculated and used to represent the variability of a splicing module across patients in said cohort. Splicing modules with a var(PSI_j_) higher than one standard deviation away of the mean var(PSI_j_) seen across patients in PENN-HTSC were deemed as “highly-variable” and characterized in downstream analyses.

### Validation of Patient Donor Cohorts

Splicing modules found in the PENN-HTSC cohort were also queried within data from an independent cohort of AML patients and a healthy myeloid donor cohort. A total of 444 RNA-seq files from AML patients were downloaded from the Beat-AML study ^11^. Moreover, patient RNA-seq files from a cohort of CD34^+^ cells isolated from 17 healthy myeloid donors were downloaded from Leucegene (https://leucegene.ca/). RNA-seq data was processed as described above. For Western blots and RT-PCR, new aliquots of cells from the same patient collection were thawed and viable blasts harvested for RNA (Trizol) and western blot (RIPA lysis buffer).

### RNA-seq Single Nucleotide Variant Calling

Patient RNA-seq files were subjected to base quality recalibration, a data pre-processing step that detects systematic errors made by the sequencer when it estimates the quality score of each base call (https://software.broadinstitute.org/gatk/). Recalibrated RNA-Seq reads were visualized within the Broad Institute Integrative Genomic Viewer (IGV_2.6.3). Clinically relevant genomic regions for AML-genes were provided by the Penn Center for Personalized Diagnostics (CPD). Single nucleotide variations (SNVs) for each AML patient within clinically relevant regions were called from their respective RNA-seq reads. We also screened for other known SNVs that have been previously tied to AML pathology. SNVs were required to be evidenced a sequencing depth larger than 10 reads.

### Gene Expression and Isoform Measurements

Transcript-level quantification of gene expression was estimated from the RNA-seq data using Salmon quasi-mapping algorithm (https://combine-lab.github.io/salmon/). Effective counts library sizes were computed using the edgeR package to account for composition biases between samples. Transcript quantification was normalized to gene-level reads per kilobase per million (RPKMs) and transformed to a measure of relative gene expression (log_2_RPKM) in downstream analyses. Variance of relative gene expression associated with batch effects was regressed out of our transcriptomic models using the ComBat empirical Bayes framework (https://rdrr.io/bioc/sva/man/ComBat.html).

### Statistical Analyses

#### Splicing modules

We plotted the distribution of splicing module variances across the cohorts and assed statistically significant differences between the mean splicing module variances using Student’s t-test. A pairwise analysis between the PSI values of each “highly-variable” splicing module generated a dissimilarity matrix of Spearman’s rho correlations. To account for the unknown vector directionality of splicing modules, we used the absolute value of the Spearman’s rank correlation coefficient in our pairwise dissimilarity matrix.

Complete linkage hierarchical clustering of the Spearman rho dissimilarity matrix was used to cluster highly correlated splicing modules into a distinct group. Splicing modules grouped together via the hierarchical clustering algorithm were used to shed light into molecular co-regulation of alternative splicing. Pearson R test was used to determine the correlation coefficients between splicing modules and gene expression vectors.

### AML Cell Lines, Western blots, and RT-PCR

In vitro growth of MV411 (ATCC CRL-9591) and HL60 (ATCC CCL-240) myeloid cell lines were by standard methods, and sequencing was performed as described in the previous method section. Radio-labeled RT-PCR was performed as previously described ^16^. Splicing modules were amplified and validated experimentally using the primer pairs specified in **Table S4**. Antibodies used for Western blots were as follows: EZH2 (a kind gift from Dr. Roberto Bonasio, UPenn), U2AF35 (Abcam catalog ab86305), and also ZRSR2 (Novus NBP1-57307).

